# Analysis of SARS-CoV-2-controlled autophagy reveals spermidine, MK-2206, and niclosamide as putative antiviral therapeutics

**DOI:** 10.1101/2020.04.15.997254

**Authors:** Nils C. Gassen, Jan Papies, Thomas Bajaj, Frederik Dethloff, Jackson Emanuel, Katja Weckmann, Daniel E. Heinz, Nicolas Heinemann, Martina Lennarz, Anja Richter, Daniela Niemeyer, Victor M. Corman, Patrick Giavalisco, Christian Drosten, Marcel A. Müller

**Affiliations:** Department of Psychiatry and Psychotherapy, University of Bonn, 53127 Bonn, Germany; Institute of Virology, Charité-Universitätsmedizin Berlin, corporate member of Freie Universität Berlin, Humboldt-Universität zu Berlin, and Berlin Institute of Health, 10117 Berlin, Germany; German Centre for Infection Research (DZIF), partner site Charité, 10117 Berlin, Germany; Max Planck Institute for Aging, 50931 Cologne, Germany; Martsinovsky Institute of Medical Parasitology, Tropical and Vector Borne Diseases, Sechenov University, 119991 Moscow, Russia

## Abstract

Severe acute respiratory syndrome coronavirus 2 (SARS-CoV-2) poses an acute threat to public health and the world economy, especially because no approved specific drugs or vaccines are available. Pharmacological modulation of metabolism-dependent cellular pathways such as autophagy reduced propagation of highly pathogenic Middle East respiratory syndrome (MERS)-CoV.

Here we show that SARS-CoV-2 infection limits autophagy by interfering with multiple metabolic pathways and that compound-driven interventions aimed at autophagy induction reduce SARS-CoV-2 propagation *in vitro*. In-depth analyses of autophagy signaling and metabolomics indicate that SARS-CoV-2 reduces glycolysis and protein translation by limiting activation of AMP-protein activated kinase (AMPK) and mammalian target of rapamycin complex 1 (mTORC1). Infection also downregulates autophagy-inducing spermidine, and facilitates AKT1/SKP2-dependent degradation of autophagy-initiating Beclin-1 (BECN1). Targeting of these pathways by exogenous administration of spermidine, AKT inhibitor MK-2206, and the Beclin-1 stabilizing, antihelminthic drug niclosamide inhibited SARS-CoV-2 propagation by 85, 88, and >99%, respectively. In sum, SARS-CoV-2 infection causally diminishes autophagy. A clinically approved and well-tolerated autophagy-inducing compound shows potential for evaluation as a treatment against SARS-CoV-2.

---

The current pandemic of severe acute respiratory syndrome coronavirus 2 (SARS-CoV-2) poses an imminent threat to global health. As of 15 April 2020, 1,878,489 individuals were infected in >200 countries, with >119,000 fatalities (*1*). SARS-CoV-2 infections cause CoV-associated disease 19 (COVID-19), which can lead to severe atypical pneumonia in humans (*2*). Currently, there are no approved therapeutics or vaccines available. The development and licensing of new FDA-approved drugs takes years, which is problematic given the urgent need for effective therapies against novel, rapidly emergent diseases like COVID-19. Antiviral drug screenings are commonly based on testing FDA-approved compound libraries against cellular and viral components (*3*). However, these undirected approaches lack functional insights into how the drugs affect virus propagation. Risk evaluations for drug repurposing and development of new therapeutics would benefit from rational drug design founded on known SARS-CoV-2-host interactions.

Compound-based targeting of cellular proteins that are essential for the virus life cycle has led to the discovery of broadly reactive drugs against a range of CoVs (*3-6*). As virus propagation strongly depends on energy and catabolic substrates of host cells, drug target identification should consider the metabolism of infected cells (*3*). Autophagy, a highly conserved cytosolic degradation process of long-lived proteins, lipids, and organelles in eukaryotic cells, is tightly controlled by metabolism (*7, 8*). During autophagy, intracellular macromolecules are recycled by incorporation into LC3B-lipidated autophagosomes (AP) and degradation into their monomers, such as fatty and amino acids, after fusion with low pH lysosomes (*9*). In the case of highly pathogenic Middle East respiratory syndrome (MERS)-CoV, we recently showed that autophagy is limited by a virus-induced AKT1-dependent activation of the E3-ligase S-phase kinase-associated protein 2 (SKP2), which targets the key autophagy initiating protein Beclin-1 (BECN1) for proteasomal degradation (*10*). Congruently, inhibition of SKP2 by different compounds, including clinically approved drugs, stabilized BECN1 and limited MERS-CoV propagation, indicating that autophagy-inducing compounds hold promise for evaluation as antiviral drugs. This paper investigates the impact of SARS-CoV-2 infection on cell metabolism and the downstream effects on autophagy, thereby identifying multiple targets for the application of approved drugs and the development of new antiviral therapies.

We aimed to characterize the effect of SARS-CoV-2 infection on autophagy by applying previously established assays (*10*) and additionally including detailed analyses of upstream autophagy regulators according to expert-curated guidelines for detecting autophagy (*11*). First, we explored whether SARS-CoV-2 reduces autophagic flux, a measure of autophagic degradation activity. We infected human bronchial epithelial cells NCI-H1299 and monkey kidney cells (VeroFM) with SARS-CoV-2 strain Munich (multiplicity of infection (MOI) of 0.0005) or heat-inactivated virus (mock-infected cells) and tested autophagic flux by co-treatment with bafilomycin A1 (BafA1). BafA1 is a specific inhibitor of vacuolar H^+^-ATPase interfering with lysosome acidification and degradation of AP cargo and preventing the fusion of APs with lysosomes to autophagy-active autophagolysosomes (AL) (*11*). Based on our previous experience (*10*), we chose a low MOI of 0.0005 and time points 8, 24, and 48 hours post infection to monitor autophagy during the exponential growth of SARS-CoV-2 reaching maximum titers of 10^7^ genome equivalents per ml (GE/ml; NCI-H1299) and 10^10^ GE/ml (VeroFM) at 48 hours post infection (**Figure S1a**). We found that 100 nM of BafA1 was sufficient to induce maximal LC3B lipidation levels in both cell cultures (**Figure S1b-d**). Immunoblotting of LC3B-II/I in the presence of 100 nM BafA1 indicated that mock-infected compared to SARS-CoV-2-infected cells showed a significant increase of LC3B-II over LC3B-I due to the BafA1-induced inhibition of autophagic flux (**Figures 1a-b**). At 8 hours post infection, LC3B-II levels were slightly elevated in SARS-CoV-2-infected cells but could still be enhanced by BafA1 treatment. At 24 and 48 hours, (vehicle- or) BafA1-treated and SARS-CoV-2 infected cells showed equally limited autophagic flux (**Figures 1a-b**). A reduction of autophagic flux was further corroborated by elevated levels of the autophagy receptor SQSTM1/P62 in SARS-CoV-2-infected cells (**Figures 1c-d**). Furthermore, we used a well-established autophagy reporter plasmid, ptfLC3, to assess lysosomal-autophagosomal fusion, a hallmark of autophagic flux (*12*). The number of ALs (in red) in mRFP and EGFP dual-tagged LC3B-expressing and SARS-CoV-2 infected NCI-H1299 cells was clearly reduced compared to mock-infected cells (**Figures 1e-f**), indicating virus-induced reduction of AP and AL fusion. Quantification showed that the total number of ALs was reduced from a mean of 88 to 62 vesicles per cell, whereas the number of APs per cell was comparable (Mock = 56 vesicles/cell; SARS-CoV-2 = 51 vesicles/cell). The ratio of APs to ALs shifted in SARS-CoV-2-infected cells, indicating strong AP accumulation (Mock= 0.707±0.053; SARS-CoV-2= 0.921 ± 0.058) (**Figure 1e**). Taken together, SARS-CoV-2 strongly reduced the autophagic flux in both cell lines, in a fashion similar to MERS-CoV (*10*).

**Fig.1:**
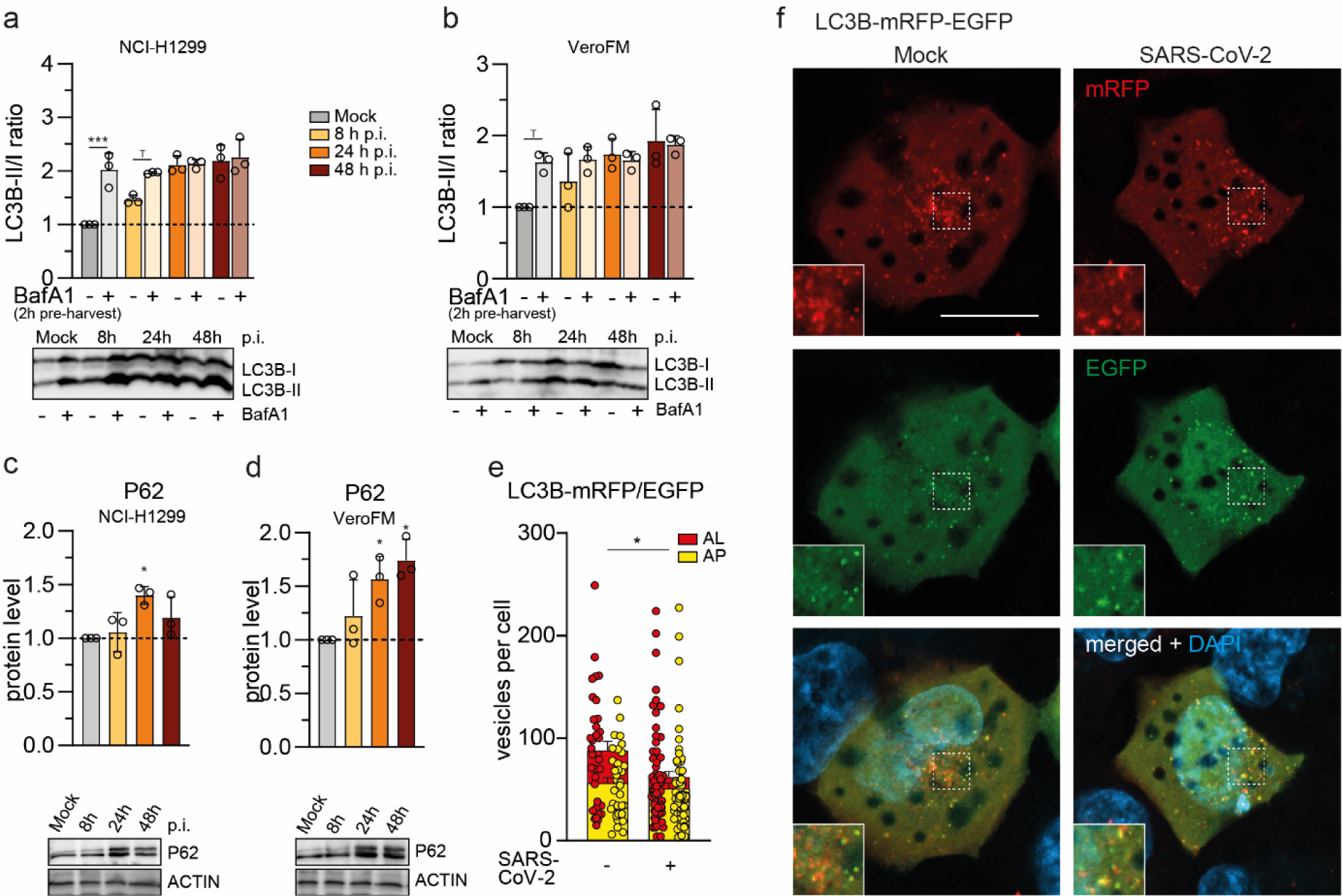
SARS-CoV-2 decreases lysosomal-autophagosomal fusion. (**a**,**b**) SARS-CoV-2 blocks autophagic flux. NCI-H1299 cells (**a**) and VeroFM cells (**b**) were infected with SARS-CoV-2 (MOI = 0.0005) and incubated with bafilomycin A1 (BafA1) or vehicle (DMSO) for 2 h before samples were harvested at 8 h, 24 h, or 48 h post infection (h p.i.). The ratios of LC3B-II/I were determined by Western blotting. (**c-d**) SARS-CoV-2 stabilizes autophagy receptor P62 protein levels. Cells were infected analogously to (a-b) and harvested at indicated time points p.i. without additional treatment. P62 protein levels were determined by Western blotting. (**e**,**f**) SARS-CoV-2 blocks fusion of autophagosomes (AP) with lysosomes. NCI-H1299 cells were transfected with tandem fluorescent-tagged LC3B (mRFP and EGFP) and infected with SARS-CoV-2 (MOI = 0.0005). Twenty-four hours later, cells were fixed and analyzed by fluorescence microscopy. Vesicles with both green and red fluorescence (APs) and with red fluorescence only (autolysosomes, AL) were counted. In all panels error bars denote standard error of mean derived from n = 3 biologically independent experiments. Tp < 0.1, *p < 0.05, ***p < 0.001 (two-way ANOVA in a,b, one-way ANOVA in c,d, t-test in e. Abbreviations: LC3B, microtubule-associated protein 1A/1B light chain 3B; mRFP, monomeric red fluorescent protein; EGFP, enhanced green fluorescent protein.

The initiation of autophagy is a cumulative result of multiple protein and signaling cascades that are involved in the energy and nutrient sensing of cells (*7, 11*). The AMP-activated protein kinase (AMPK) and the mammalian target of rapamycin complex 1 (mTORC1) kinase are in continuous crosstalk with glucose and protein homeostasis respectively (*13, 14*). We monitored key proteins of the AMPK and mTORC1 pathways and their regulation/degradation by Western blot analyses after infection with SARS-CoV-2 (**Figure 2**). SARS-CoV-2 infection influenced most of the analyzed components of the AMPK/mTORC1 pathway. Phosphorylated, active forms of AMPK, AMPK substrates (LXRXX), AMPK downstream targets (TSC2 and ULK1), and mTORC1 were downregulated upon SARS-CoV-2 infection, suggesting a virus-induced reduction of cellular glycolysis, protein translation, and cell growth. In addition, we found increased levels of phosphorylated AKT1, which activates the negative regulator of BECN1, SKP2 (*10*). Low BECN1 levels and subsequently reduced ATG14 phosphorylation and oligomerization (**Figure 2, lower left panel**) explain the lack of fusion of APs with lysosomes observed in **Figure 1e-f** as ATG14 oligomers facilitate fusion via SNARE-proteins, SNAP29 and STX17 (*10*). In summary, SARS-CoV-2 alters autophagy-relevant signaling and at the same time appears to hamper AP/lysosome fusion efficiency via autophagy.

**Fig.2:**
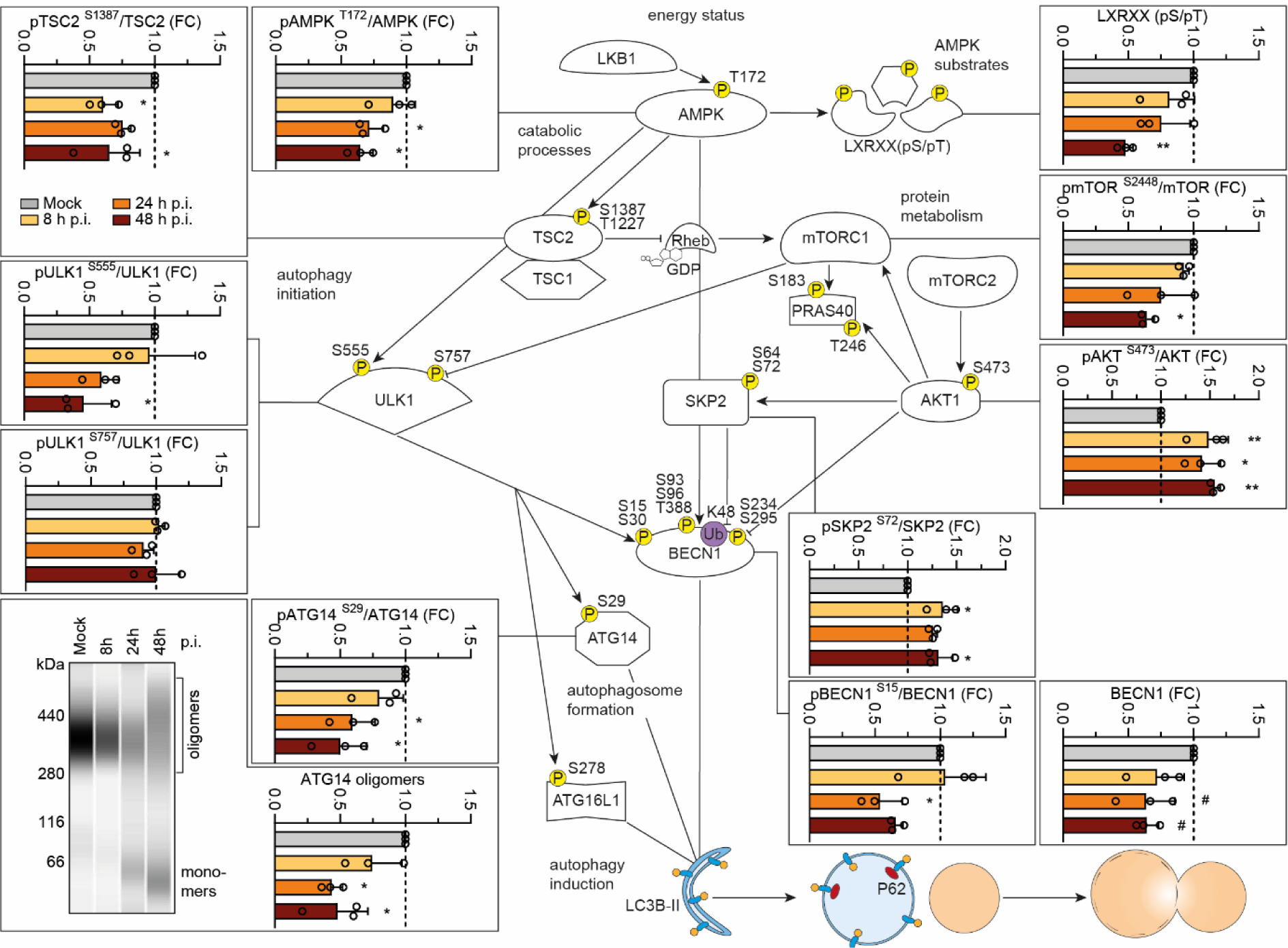
SARS-CoV-2 hijacks autophagy signaling on multiple regulatory levels. VeroFM cells were infected with SARS-CoV-2 (MOI = 0.0005) and harvested at 8 h, 24 h, or 48 h post infection (h p.i.). Protein levels and phosphorylation status of selected autophagy-relevant proteins were determined by Western blotting. For analysis of ATG14 oligomers (bottom panel, left) cells were incubated with a chemical crosslinker (DSS) 2 h prior to cell harvest (for detailed protocol see methods section). Cell extracts were separated and immunoblotted using Wes (ProteinSimple) capillary electrophoresis. Error bars in all panels denote standard error of mean derived from n = 3 biologically independent experiments. Tp < 0.1, *p < 0.05, ***p < 0.001 (one-way ANOVA; Bonferroni correction (post hoc)). Abbreviations: AMPK, AMP-activated protein kinase; TSC1/2, tuberous sclerosis 1/2; mTORC1/2, mammalian target of rapamycin complex 1/2; PRAS40, proline-rich AKT1 substrate 1; SKP2, S-phase kinase-associated protein 2; BECN1, Beclin-1; ATG, autophagy-related; Rheb, Ras homolog enriched in brain; ULK1, Unc-51-like kinase 1;

To further explore the direct impact of SARS-CoV-2 infection on AMPK/mTORC1, we performed a metabolomics profiling analysis in VeroFM cells 24 hours post infection using an elevated MOI of 0.1 to guarantee ubiquitous infection of cells (**Figure 3**). The metabolite profiles were analyzed by multivariate principal component analysis (PCA), an unsupervised statistical method suitable for analyzing and classifying metabolomics datasets (*15*). The profiles clearly separated into SARS-CoV-2 and control group (**Figure S2a**). Altogether, the levels of 25 metabolites were significantly altered by SARS-CoV-2 infection (**Supplementary Table 1**). We performed pathway analyses in order to gain insights into the cellular and mechanistic consequences of a SARS-CoV-2 infection. We retrieved eight affected pathways using the significantly altered metabolites from **Supplementary Table 1** (**Figure 3a**). Glutathione (GSH) metabolism, pyrimidine metabolism, and aminoacyl-tRNA biosynthesis had the most pronounced changes. Metabolites of the butanoate metabolism as well as the alanine, aspartate and glutamate metabolism were also elevated. In contrast, the tricarboxylic acid (TCA) cycle, glutamine and glutamate metabolism, and glyoxylate and dicarboxylate metabolism were reduced by SARS-CoV-2. The volcano plot visualizes the metabolites with large magnitude fold changes. We identified seven altered metabolites: Cys-glycine, N-acetylputrescine, adenosine, putrescine, homocysteine and cysteine levels were elevated whereas fructose-6P was strongly depleted (**Figure 3b**). Putative links between the different pathways during SARS-CoV-2 infection are shown in **Figure 3c**. We observed elevated ATP levels that might facilitate energy-dependent processes such as virus replication and entry (**Figure 3c, blue**). High lactic acid and low fructose-6P levels indicate exhaustive glycolysis activity (**Figure 3c, green**). Decreased levels of TCA metabolites (**Figure 3c, purple**) further support a switch from (mitochondria-dependent) oxidative phosphorylation to lactic acid fermentation, to provide NAD+ for glycolytic ATP production. Consequently, we observed a reduced AMP/ATP ratio (**Figure 3c, blue**) limiting AMPK activation (*16*) and subsequently autophagy (see **Figure 2**). Limited autophagy might prevent degradation of viral products and might enhance the availability of non-autophagic membrane material to generate double-membrane vesicles that are generated during coronavirus replication (*17, 18*). Interestingly, despite reduction of protein-degrading autophagy, we observed strong upregulation of amino acids, especially those linked to glutathione metabolism, which is responsible for elimination of reactive oxygen species (ROS). This might be essential to limit the mitochondrion-derived ROS that are generated during excessive ATP production (*19*) and should be investigated in future studies. Further investigations should also clarify why the observed upregulated amino acid levels do not activate mTORC1 (see **Figure 2**, pmTOR^S2448^/mTOR and pULK1^S757^/ULK1). However, spatially distinct pools of mTORC1 proteins might serve as an explanation (*20*).

**Fig.3:**
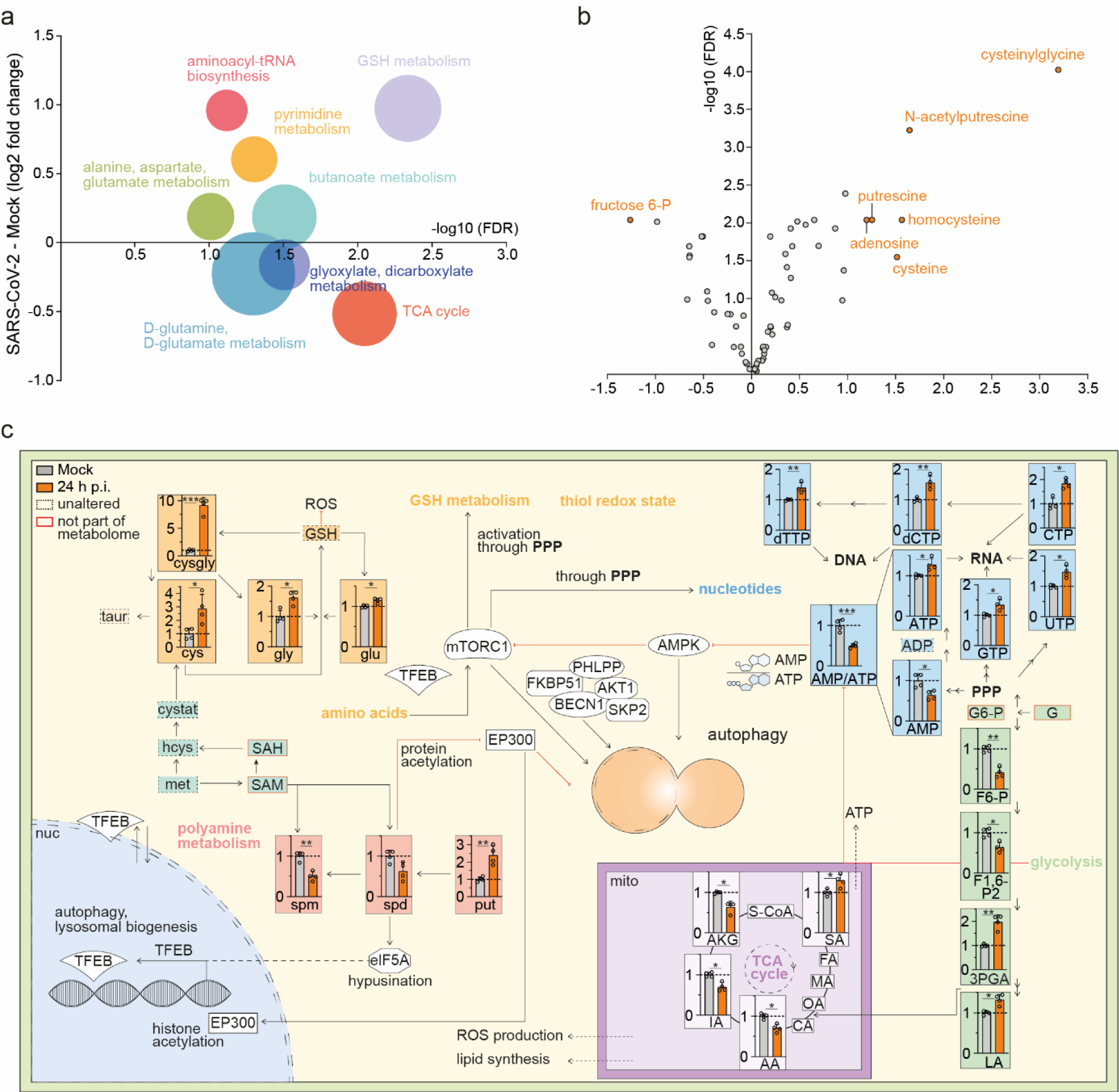
SARS-CoV-2 affects key metabolic pathways. (**a**) Analysis and regulation of significantly altered pathways of mock and SARS-CoV-2 infected (24 h p.i) VeroFM cells. The f(x)-axis shows the (median) log2 fold change (FC) of all significantly altered metabolites of the indicated pathway and the –log10 adjusted p-value (false discovery rate (FDR)) is shown on the x-axis. The size of the circles illustrates the number of significantly changed metabolites in relation to all metabolites of a specific pathway. N = 4 per group. (**b**) Volcano plot of metabolome of SARS-CoV-2 infected (24 h.p.i) VeroFM cells. Metabolites with log2(FC) ≥ 2 and –log10(FDR) ≥ 1.3 were considered significant. N = 4 per group. (**c**) Analysis of the autophagic pathway and the involved metabolites: ‘amino acids’ and ‘GSH metabolism’ (orange), ‘nucleotides’ (blue), ‘glycolysis/ TCA cycle’ (violet) and ‘polyamine metabolism’ (red) and ‘AMP/ATP ratio’ (blue) upon mock and SARS-CoV-2 infected (24 h p.i) VeroFM cells. Error bars represent standard deviations *Adjusted p-value (FDR) ≤ 0.05; **FDR ≤ 0.01; ***FDR ≤ 0.001. N=4 per group. Abbreviations: taur, taurine; cysgly, cysteinylglycine; cys, cysteine; gly, glycine; glu, glutamate; cystat, cystathionine; hcys, homocysteine; met, methionine; SAH, S-adenosyl-L-homocysteine; SAM, S-adenosylmethionine; ROS, reactive oxygen species; GSH, glutathione; spm, spermine; spd, spermidine; put, putrescine; S-CoA, Succinyl-CoA; SA, succinic acid; FA, fumaric acid; MA, malic acid; CA, citric acid; AA, aconitic acid; IA, isocitric acid; AKG, α-ketoglutaric acid; LA, lactic acid; 3PGA, 3-phosphoglyceric acid; F1,6-P2, fructose 1,6-bisphosphate; F6-P, fructose 6-phosphate; G, glucose; G6-P, glucose 6-phosphate; AMP, adenosine monophosphate; ADP, adenosine diphosphate; ATP, adenosine triphosphate; GTP, guanosine triphosphate; UTP, uridine triphosphate; CTP, cytidine triphosphate; dCTP, deoxycytidine triphosphate; dTTP, deoxythymidine triphosphate; PPP, pentose phosphate pathway; TCA, tricarboxylic acid; FKBP51, 51 kDa FK506-binding protein; EP300, histone acetyltransferase p300; TFEB, transcription factor EB; elF5A, eukaryotic translation initiation factor 5A; PHLPP, PH domain leucine-rich repeat-containing protein phosphatase; nuc, nucleus; mito, mitochondrion.

Interestingly, putrescine (**Figure 3c, red**) was increased whereas its downstream products of the polyamine biosynthesis pathway, spermidine and spermine, were strongly reduced. The putrescine increase might be explained by a SARS-CoV-2-induced inhibition of spermidine synthase, the enzyme that synthesizes spermidine from putrescine (**Figure 3c, red**). Spermidine is an important polyamine capable of inducing autophagy (*21*) and responsible for hypusination of the eukaryotic translation initiation factor eIF5A (*22*). eIF5A activates the autophagy transcription factor TFEB and regulates proteins responsible for mitochondrial respiration (*23*).

Next, we targeted different components of the metabolic/autophagic pathway by exogenous administration of selective inhibitors, approved drugs, or cell metabolites, and explored the effect on SARS-CoV-2 propagation (**Figure 4**). Cell viability tests were performed for inhibitors and drugs to exclude toxicity (**Figure S3a**). VeroFM cells were infected with an MOI of 0.0005 and treated 1 hour post infection with different concentrations of each substance according to the manual instructions or previous publications. Virus growth was monitored by real-time RT-PCR in cell culture supernatants 24 and 48 hours post infection (**Figure S3b-d**). DFMO (500 µM) blocks ornithine-to-putrescine synthesis (*24*) and was previously described as an RNA virus inhibitor upon pretreatment (*25*) but had minor effects on SARS-CoV-2 growth upon post-treatment, whereas spermidine (100 µM) and spermine (100 µM) inhibited SARS-CoV-2 propagation by up to 66% (**Figure 4a, upper left panel, Figure S3b**,**c**). AICAR (25 µM), a known AMPK activator (*26*), slightly induced virus growth at 24 hours post infection (**Figure 4a, upper right panel**), which seems contradictory, as AMPK is downregulated upon SARS-CoV-2 infection (see **Figure 2**), but could be explained by AMPK-induced mTORC1 inhibition or AMPK-unrelated functions of AICAR (*27*). Alternatively, AICAR-dependent AMPK activation might be insufficient to restore autophagic flux (see **Figure 1**). In addition, mTORC1 inhibition by rapamycin (0.3 µM) clearly induced virus growth (**Figure 4a, right panel, Figure S3d**,**e**). Rapamycin, which has previously been proposed as a treatment option for COVID-19 (*28*), possibly enhances the inhibitory effects of SARS-CoV-2 on mTORC1 (see **Figure 2**) to reduce cellular protein translation, which would serve as an additional explanation for the above-mentioned elevated amino acid levels (see **Figure 3c, orange**). These results suggest that detailed molecular and functional analyses should be performed for rapamycin before considering its use in clinical trials. The AKT1 inhibitor MK-2206 (1 µM), which is currently being tested in a clinical phase 2 study against breast cancer (*29*), reduced SARS-CoV-2 propagation up to 50% (**Figure 4a, lower right, Figure S3d**,**e**). AKT1 blocks mTORC1 inhibitor TSC2 (*30*) and further supports the suggestion that up-regulation of mTORC1 components has antiviral effects. As AKT1 inhibition results in BECN1 up-regulation and autophagy induction (*10, 31*), SARS-CoV-2 growth inhibition was expected. Direct blocking of the negative BECN1 regulator SPK2 by previously described inhibitors SMIP004, SMIP004-7, valinomycin, and niclosamide (*10*) showed SARS-CoV-2 growth inhibition from 50 (SMIP004, SMIP004-7) to over 99% in case of valinomycin and niclosamide (**Figure 4a, lower panel, Figure S3d**,**e**). We further confirmed that the dominant intervention of niclosamide during SARS-CoV-2 infection acts on autophagy induction, as adding BafA1 after niclosamide treatment showed an enhancing effect on the lipidation of LC3B as reflected by comparable LC3B-II/I ratios between mock- and SARS-CoV-2-infected cells (**Figure 4b**). However, we cannot exclude that the activity of niclosamide as a hydrogen ionophore has additional inhibitory functions, e.g. by blocking endosomal acidification (*32*), which is important for SARS-CoV-2 entry (*6*). We further used our in vitro model to explore the possibility of prophylactic treatment with spermidine and niclosamide, which are well-tolerated, clinically applied (*33*) or FDA-approved (*34*) compounds, respectively. To assess the efficacy of pretreatment, VeroFM cells were preincubated with spermidine (100 μM) or niclosamide (5 μM) for 24 hours. Cells were infected with SARS-CoV-2 using an MOI of 0.05 without further compound treatment and virus growth was monitored in supernatants for 24 hours (**Figure 4c, Figure S3f**). In both cases, SARS-CoV-2 growth was reduced by 70%, suggesting that both compounds exhibit long-lasting antiviral effects, and supporting further investigation into prophylactic use of these compounds. For clinical use, half-maximal inhibitory concentration (IC50) of compounds should be in a non-toxic range and reach adequate plasma levels (*35*). The IC50 was 149 μM (R^2^ = 0.71) for spermidine, 0.09 μM (R^2^= 0.95) for MK-2206, and 0.17 μM (R^2^= 0.63) for niclosamide, based on plaque-forming infectious units (**Figure 4c)**. Maximal inhibition of infectious virus at non-toxic concentrations was identified at 333.3 μM for spermidine (85%), at 3.7 μM for MK-2206 (88%), and at 1.24 μM for niclosamide (>99%). Whereas plasma levels and pharmacokinetics of spermidine are yet to be evaluated, peak plasma concentrations were 0,176 μg/ml (0.43 μM) for MK-2206 (*36*) and 0.25 to 6.0 μg/ml (0.76-18.35 μM) for niclosamide (*37*), encouraging use in clinical trials.

**Fig.4:**
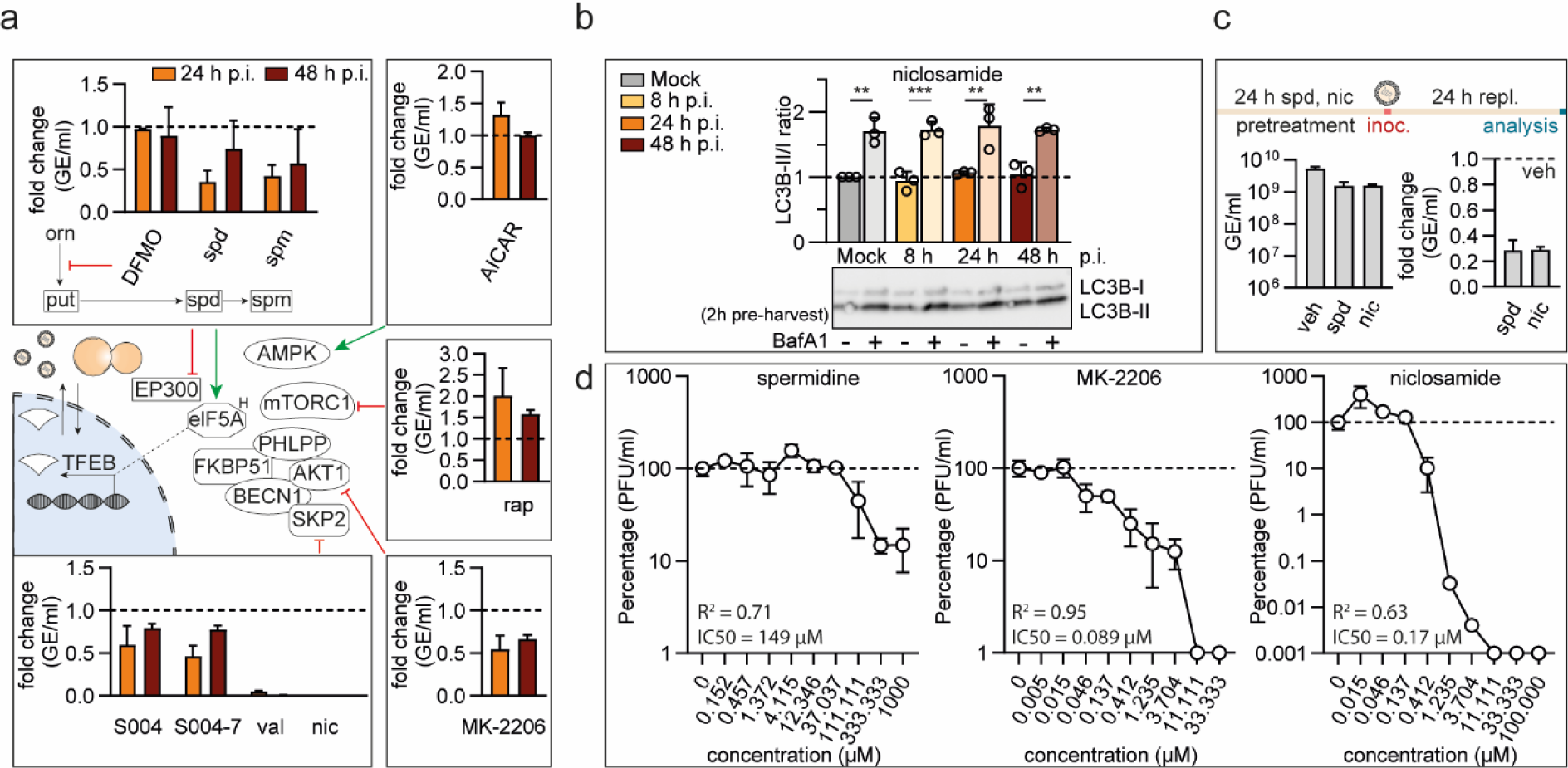
BECN1-stabilizing compounds and polyamines inhibit SARS-CoV-2 growth in cell cultures. (**a**) Schematic representation of autophagy signaling indicating site of action of small-molecule inhibitors used for pathway modulation, tested in SARS-CoV-2 growth assays. VeroFM cells were infected with SARS-CoV-2 (MOI = 0.0005) and treated with (top, left) DFMO (500 µM), spd (spermidine, 100 µM), spm (spermine, 100 µM), (top right) AICAR (25 µM), (middle, right) rap (rapamycin, 300 nM), (bottom, left) SKP2 inhibitors S004 (SMIP004, 10 µM), S004-7 (SMIP004-7, 10 µM), val (valinomycin, 5 µM), nic (niclosamide, 10 µM), (bottom, right) MK-2206 (1 µM) or DMSO (vehicle, dashed lines). SARS-CoV-2 genome equivalents per mL (GE) were determined by real-time RT-PCR at 24 h and 48 h p.i., data are presented as fold difference. (**b**) Niclosamide treated (10 µM) VeroFM cells were infected with SARS-CoV-2 (MOI = 0.0005) and incubated with bafilomycin A1 (BafA1, 100 nM) or vehicle (DMSO) for 2 h before samples were harvested at 8 h, 24 h, or 48 h post infection (h p.i.). The ratios of LC3B-II/I were determined by Western blotting. (**c**) VeroFM cells were treated with spd (100 µM), nic (5 µM) or veh (vehicle) 24 h prior to infection with SARS-CoV-2 (MOI = 0.05). SARS-CoV-2 genome equivalents per ml (GE) were determined by real-time RT-PCR at 24 h p.i., data are presented GE/mL (left) or as fold difference (right). (**d**) Concentration-dependent inhibition of SARS-CoV-2 growth by spermidine, MK-2206, and niclosamide. VeroFM cells were infected with SARS-CoV-2 (MOI = 0.0005) and treated with different concentrations of spermidine, MK-2206, and niclosamide as shown in the figure. SARS-CoV-2 plaque forming units (PFU) were determined at 24 h (spermidine, MK-2206) and 48 h (niclosamide) p.i. by plaque assay. Data are presented as virus growth in percent. In all panels error bars denote standard error of mean derived from n = 3 biologically independent experiments. Abbreviations: orn, ornithine; DFMO, difluoromethylornithine; elF5AH, hypusinated elF5A; PFU, plaque-forming unit.

In summary, our data show that highly pathogenic SARS-CoV-2 reprograms the metabolism of cells and limits AMPK/mTORC1 activation and autophagy. Our mechanistic approach, including metabolism, protein/phosphoprotein analyses and targeted SARS-CoV-2 inhibition assays, identified multiple cellular targets for the development of new and application of already available antiviral compounds.

## Acknowledgments

We thank Patricia Tscheak and Antje Richter (Charité) for excellent technical assistance and Terry Jones for editing of the manuscript.

## Author contributions

N.C.G. and M.A.M. designed and conceived the work. N.C.G., J.P., T.B., F.D., J.E., N.H., D.E.H., M.L., A.R., M.A.M. carried out experiments. N.C.G., J.P., T.B., J E., K.W. V.M.C., D.N., P.G., C.D., M.A.M. analyzed data or contributed essential material. N.C.G., M.A.M. wrote the main paper text. N.C.G., T.B. prepared all figures. All authors reviewed the paper.

