## Supplementary Material for "Analysis of SARS-CoV-2-controlled autophagy reveals spermidine, MK-2206, and niclosamide as putative antiviral therapeutics"

### SUPPLEMENTARY MATERIAL Gassen et al.

### Material & Methods, Supplementary Figures, Supplementary Tables

#### Material and Methods

##### *Chemicals*

The following chemicals were used for treatment of cells: BafA1 (Alfa Aeser, J61835), RAP (Tocris, 1292), NIC (Sigma-Aldrich, Munich, Germany, N3510), VAL (Sigma-Aldrich, V0627), MK-2206 HCl (Cayman Chemical, 11593), SMIP004 (Maybridge Chemicals, TL00829SC), SMIP004-7 (Maybridge Chemicals, SB01607DA), DSS (disuccinimidyl suberate, Thermo Fisher Scientific, 21655), Spermine (Sigma-Aldrich, S4264-5G), Spermidine (Sigma-Aldrich, 85558-5G), DFMO (Sigma-Aldrich, D193-25MG).

##### *Cells*

VeroFM (ATCC CCL-81), VeroE6 (ATCC CRL-1586), and NCI-H1299 (ATCC CRL-5803) were cultivated in Dulbecco's Modified Eagle´s Medium (DMEM) supplemented with 10% fetal bovine serum (FBS), 1% penicillin/streptomycin, 1% non-essential amino acids, and 1% sodium pyruvate at 37 °C and 5% $CO_2$.

##### *Virus strains and infection*

For virus infection with SARS-CoV-2 strain Munich 984, 2 x 10e5 cells ml-1 were seeded in 6-well plates. After 24 h, cells were infected with an MOI = 0.0005 in serum-free medium. In parallel, mock infected cells were inoculated with heat-inactivated (95 °C, 10 min) virus. After 1 hour, the virus dilutions were removed and the wells were washed twice with PBS and refilled with DMEM (supplemented as described above). Samples were taken at the indicated time points (8, 24, and 48 hours). All virus infection experiments were conducted under biosafety level 3 conditions with enhanced respiratory personal protection equipment. For viral RNA extraction, 50 µl of cell culture supernatant was mixed with 300 µl MagNA Pure external lysis buffer (Roche, Penzberg, Germany) followed by 70 °C for 10 minutes. Extraction was done by automated pipetting using MagNA Pure 96 instrument (Roche).

##### *Real-time reverse-transcription PCR*

SARS-CoV-2 genome equivalents were detected by real-time RT-PCR assay targeting SARS-CoV-2 E gene as reported before (*1*), using the following primers: E_Sarbeco_F: ACAGGTACGTTAATAGTTAATAGCGT; E_Sarbeco_P1: FAM-ACACTAGCCATCCTTACTGCGCTTCG-BBQ; E_Sarbeco_R: ATATTGCAGCAGTACGCACACA. Briefly, real-time RT-PCRs were performed using the Superscript III OneStep RT-PCR kit (Invitrogen, Darmstadt, Germany). Reactions were performed in 25 µl volume with 12.5 µl of 2x reaction buffer, 2 µl of 10 µM forward and reverse primers, 1 µl of 10 µM probe, 1 µl of bovine serum albumin (BSA), 1 µl enzyme mix, RNase-free water, and 5 µl of RNA template. RNA was reverse transcribed at 55°C for 10 min, followed by initial denaturation at 95°C for 180 s. Cycling and fluorescence signal acquisition was performed for 45 cycles of 95 °C for 15 s and 58 °C for 30 s. Real-time RT-PCR experiment and data processing was done using the LightCycler® 480 Real-Time PCR System (Roche). Absolute quantification was performed using SARS-CoV-2-specific in vitro-transcribed RNA standards, as described before.

*Plaque assay*

Infectious SARS-CoV-2 plaque forming units (PFU) were quantified by plaque titration on VeroE6 cells, as described previously (*2*), with minor modifications. Briefly, VeroE6 monolayers were seeded in 24-well plates, washed with PBS, incubated with serial dilutions of SARS-CoV-2-containing cell culture supernatants in duplicates, and overlaid with 1.2% Avicel in DMEM, supplemented as described above. After 72 h, cells were fixated with 6% formaline and visualized by crystal violet staining. Subsequently, plaques were counted and PFU/ml were determined.

*Metabolite extraction for Liquid Chromatography mass spectrometry (LC-MS)*

VeroFM cells (5x10e6 cell/ml) were seeded into 6-well (quadruplicates) and infected with SARS-CoV-2 or heat-inactivated SARS-CoV-2 (mock) at an MOI of 0.1 as described above. After 24 hours, supernatants were discarded and the adherent cells were washed twice with 1 ml ice-cold washing buffer containing 75 mM ammonium carbonate (pH7.4). Subsequently, metabolite extraction from each cell culture was performed using 400 µl of a mixture of 40:40:20 [v:v:v] of pre-chilled (-20°C) acetonitrile:methanol:water (Optima™ LC/MS grade, Thermo Fisher Scientific). Supernatant was collected in fresh 1.5 ml tubes. Remaining cells were extracted a second time with 400 µl of the extraction mix and cells were carefully scratched from the plate surface. Extracts were combined and heated to 70°C for 10 min, to inactivate virus, before centrifuging them for 10 min at 21,100 x g. The metabolite-containing supernatant was collected in fresh tubes and stored at -80°C. Supernatants were concentrated to dryness in a Speed Vac concentrator (Eppendorf).

*LC-MS analysis of amine-containing metabolites*

For amino acid analysis the benzoylchlorid derivatization method (*3*) was used. In brief: the dried metabolite pellets were resuspended in 90 µl of the LC-MS-grade water (Milli-Q 7000 equipped with an LC-Pak and a Millipak filter, Millipore). 20 µl of the re-suspended sample was mixed with 10 µl of 100 mM sodium carbonate (Sigma-Aldrich) followed by the addition of 10 µl 2% benzoylchloride (Sigma-Aldrich) in acetonitrile (Optima-Grade, Fisher-Scientific). Samples were vortexed before centrifugation for 10 min at 21,300 x g at 20°C. Clear supernatants were diluted 1:10 with LC MS-grade water and transferred to fresh auto sampler tubes with conical glass inserts (Chromatographie Zubehoer Trott) and analyzed using a Vanquish UHPLC (Thermo) connected to a Q-Exactive HF (Thermo).

For the analysis 1 µl of the derivatized sample were injected onto a 100 x 2.1 mm HSS T3 UPLC column (Waters). The flow rate was set to 400 µL/min using a buffer system consisted of buffer A (10 mM ammonium formate (Sigma-Aldrich), 0.15% formic acid (Sigma-Aldrich) in LC MS-grade water) and buffer B (acetonitrile, Optima-grade, Fisher-Scientific). The LC gradient was: 0% B at 0 min; 0-15% B 0-0.1 min; 15-17% B 0.1-0.5 min; 17-55% B 0.5-7 min, 55-70% B 7-7.5 min; 70-100% B 7.5-9 min; 100% B 9-10 min; 100-0% B 10-10.1 min, 10.1-15 min 0% B. The mass spectrometer was operating in positive ionization mode monitoring the mass range m/z 50-750. The heated ESI source settings of the mass spectrometer were: Spray voltage 3.5 kV, capillary temperature 250°C, sheath gas flow 60 AU and aux gas flow 20 AU at a temperature of 250°C. The S-lens was set to a value of 60 AU.

Data analysis was performed using the TraceFinder software (Version 4.2, Thermo Fisher Scientific). Identity of each compound was validated by authentic reference compounds, which were analysed independently. Peak areas were analysed by using extracted ion chromatogram (XIC) of compound-specific $[M + nBz + H]^+$ where n corresponds to the number of amine moieties which can be derivatized with a bezoylchlorid (Bz). XIC peaks were extracted with a mass accuracy (<5 ppm) and a retention time (RT) tolerance of 0.2 min.

*Anion-Exchange Chromatography Mass Spectrometry (AEX-MS) of the analysis of TCA cycle and glycolysis metabolites*

Anion-Exchange Chromatography was performed simultaneously to the LC MS analysis. 50 µl of the re-suspended sample were diluted 1:5 with LC MS-grade water and analysed using a Dionex ionchromatogrypy system (ICS 5000, Thermo Scientific). The applied protocol was adopted from (*4*). In brief: 10 µL of polar metabolite extract were injected in full loop mode using an overfill factor of 3, onto a Dionex IonPac AS11-HC column (2 mm × 250 mm, 4 µm particle size, Thermo Scientific) equipped with a Dionex IonPac AS11-HC guard column (2 mm × 50 mm, 4 µm, Thermo Scientific). The column temperature was held at 30°C, while the auto sampler was set to 6°C. A potassium hydroxide gradient was generated by the eluent generator using a potassium hydroxide cartridge that was supplied with deionized water. The metabolite separation was carried at a flow rate of 380 µL/min, applying the following gradient. 0-5 min, 10–25 mM KOH; 5-21 min, 25–35 mM KOH; 21-25 min, 35-100 mM KOH, 25-28 min, 100 mM KOH, 28-32 min, 100-10 mM KOH. The column was re-equilibrated at 10 mM for 6 min.

The eluting metabolites were detected in negative ion mode using ESI MRM (multi reaction monitoring) on a Xevo TQ (Waters) triple quadrupole mass spectrometer applying the following settings: capillary voltage 1.5 kV, desolvation temperature 550°C, desolvation gas flow 800 L/h, collision cell gas flow 0.15 mL/min. All peaks were validated using two MRM transitions one for quantification of the compound, while the second ion was used for qualification of the identity of the compound. The settings for the MRM transitions are given in **Supplementary Tables 1, 2.** Data analysis and peak integration was performed using the TargetLynx Software (Waters).

*Analysis of metabolomic data*

Identification of significant metabolite level alterations: Metabolite intensities were sample log2 transformed and pareto scaled for further statistical analysis. Significant metabolite level changes upon 24 h p.i. compared to mock were identified by student's T-test with an adjusted p-value (false discovery rate (FDR)) ≤ 0.05 using MetaboAnalyst (*5*).

Identification of significantly overrepresented pathways: Metabolomics pathway analyzes were performed with the significantly altered metabolites from Supplementary Table 1 using MetaboAnalyst applying a Fisher's Exact Test and Out-degree Centrality for pathway topology analysis. Pathways were considered for further analyses with an FDR ≤ 0.1 (*5*).

*Toxicity assays*

MTT assay. VeroFM or NCI-H1299 cells were incubated in the presence of 0.5 mg mL−1 tetrazole 3-(4,5-dimethylthiazol-2-yl-)-2,5-diphenyltetrazolium bromide (MTT) for 4 h at 37 °C and 5% $CO_2$. Read-out of MTT assay was carried out as described previously (*6*).

*Atg14 oligomerization*

The PBS-washed cell pellet was incubated with 75 µM DSS (disuccinimidyl suberate, Thermo Fisher Scientific, 21655) or corresponding vehicle (DMSO) in PBS. Crosslinking was performed for 30 min at room temperature followed by 2 h at 4 °C while rotating. After washing with PBS, crosslinking was quenched in tris-buffered saline (pH 7.0) for 20 min at 4 °C. Cells were lysed in Pierce™ IP lysis buffer and analyzed by capillary electrophoresis on Wes™ (ProteinSimple) using the 60–440 kDa cartridges.

*Western blot analysis*

Protein extracts were obtained by lysing cells in Pierce™ IP lysis buffer (150 mM NaCl, 1% NP-40, 1 mM EDTA, 5% glycerol, 25 mM Tris-HCl (pH7.4); ThermoFisher Scientific) freshly supplemented with protease inhibitor (Merck Millipore, Darmstadt, Germany), benzonase (Merck Millipore), 5 mM DTT (Sigma), and 1% PhosSTOP™ phosphatase inhibitor (Roche). Proteins were separated by SDS-PAGE and electro-transferred onto PVDF membranes. Blots were placed in Tris-buffered saline, supplemented with 0.05% Tween (Sigma Aldrich) and 5% non-fat milk for 1 h at room temperature and then incubated with primary antibody (diluted in TBS/0.05% Tween) overnight at 4 °C while shaking. The following primary antibodies were used: beta-actin (1:5,000 Cell Signaling Technology, #8457), SQSTM1/p62 (1:1,000, Cell Signaling Technology, #5114), LC3-B (1:1,000, Cell Signaling Technology, #3868), Beclin-1 (1:1,000, Cell Signaling Technology, #3738), p-Beclin1 (S15) (1:1,000, Cell Signaling Technology, #84966), Atg14 (1:1,000, Cell Signaling Technology, #5504), pAtg14 (S29) (1:1,000, Cell Signaling Technology, #13155), Ulk1 ULK1 (1:1,000, Cell Signaling Technology, #8054), pUlk1 pULK1 (S555) (1:1,000, Cell Signaling Technology, #5869), pUlk1 pULK1 (S757) (1:1,000, Cell Signaling Technology, #6888), TSC2 (1:1,000, Cell Signaling Technology, #3612), pTSC2 (S1387) (1:1,000, Cell Signaling Technology, #5584), AMPKαalpha (1:1,000, Cell Signaling Technology, #2532), pAMPK (T172) (1:1,000, Cell Signaling Technology, #2531), pAMPK substrate motif (1:1,000, Cell Signaling Technology, #5759), pAKT (S473) (1:1,000, Cell Signaling Technology, #4060), AKT (1:1,000, Cell Signaling Technology, #9272), mTOR (1:1,000, Cell Signaling Technology, #2983), p-mTOR (S2448)(1:1,000, Cell Signaling Technology, #5536), HSC70 (1:5,000, Enzo Life Sciences, ADI-SPA-757-F). Secondary antibodies: anti-rabbit IgG, HRP-linked antibody (1:10,000, Cell Signaling Technology, #7074), anti-mouse IgG, HRP-linked antibody (1:10,000, Cell Signaling Technology, #7076)

Subsequently, blots were washed and probed with the respective horseradish peroxidase- (or fluorophore-conjugated) secondary antibody for 1 h at room temperature. The immuno-reactive bands were visualized using ECL detection reagent (BioRad, Hercules, CA, USA). Determination of the band intensities were performed with BioRad, ChemiDoc MP.

In general, protein quantification was performed by normalization to the intensity of actin, which was determined on the same membrane. For quantification of lipidated LC3B, the intensity of LC3B-II was always referred to the signal intensity of the corresponding LC3B-I, following the guidelines for the determination of autophagy (*7*). For quantification of phosphorylated proteins this signal was always referred to the signal intensity of the corresponding total protein.

*Lysis of SARS-CoV-2 infected cells*

Whole cell lysates were prepared with Pierce™ IP lysis buffer (150 mM NaCl, 1% NP-40, 1 mM EDTA, 5% glycerol, 25 mM Tris-HCl (pH7.4); ThermoFisher Scientific), supplemented with protease inhibitor (Merck Millipore), benzonase (Merck Millipore), 5 mM DTT (Sigma Aldrich, Munich, Germany), and 1% PhosSTOP™ phosphatase inhibitor (Roche). For cell lysis, supernatant was discarded and cells were carefully rinsed 3x with pre-cooled 1x PBS. 100 µl of pre-cooled IP lysis buffer was added and cells were scraped off, transferred to a microcentrifuge tube, and incubated for 20 min at 4 °C for efficient lysis. Then, SDS loading buffer (4x NuPAGE LDS, ThermoFisher Scientific) was added and samples were heated to 95 °C for 10 min. Samples were stored at -80 °C and processed as indicated in the respective sections.

*Autophagic flux*

To determine the effect of drug treatment or CoV infection on autophagic flux, DMSO (vehicle control) or BafA1 were applied two hours before cell lysis, followed by Western blot analysis as described. In order to detect autophagosomes and autophagolysosomes by immunofluorescence test, VeroFM and NCI-H1299 cells were transfected with ptfLC3 plasmid (Addgene.org, #21074) that expresses LC3 tagged with both GFP (inactivated in autolysosomes) and mRFP (resists inactivation in autolysosomes) (*8*), following the guidelines for monitoring autophagy (*7*). For transfections, cells were treated with 0.5 µg plasmid and 1.5 µl Fugene HD (Promega, Mannheim, Germany) 24 hours prior to infection with SARS-CoV-2 (MOI = 0.001) and drug treatment the next day. Cells were fixed (6% formaldehyde for 1 h) at indicated time points (8 and 24 hours), glass slides were mounted with ProLong™ Gold Antifade Mountant with DAPI (ThermoFisher Scientific), and analyzed by fluorescence microscopy (AxioVert 200 M, Carl Zeiss, Oberkochen, Germany) equipped with a plan-Apochromat 63x/1.40 Oil DIC objective and an AxioCam MR R3 camera. Images were acquired and processed using AxioVision software ZEN Pro 2 (Carl Zeiss). Vesicles were counted by an experimenter blind to the conditions. Additionally, formaline-fixed cells were incubated for 1 h at 37° C with heat-inactivated human anti-SARS-CoV-2 serum (diluted 1:100) taken from a SARS-CoV-2 patient (serum collection National consiliary laboratory for CoV diagnostics at Charité, Berlin). Secondary antibody incubation was done with Alexa fluor 647-labelled polyclonal goat-anti human antibody for 45 min at 37° C (1:200, Thermo Fisher Scientific, #A-21445).

*Statistical analysis*

When two groups were compared, the student's t-test was applied. For three or more group comparisons, one or two-way analysis of variance (ANOVA) was performed, as appropriate, followed by Tukey's or Bonferroni's post hoc test, as appropriate. All t-test t- and p-values and ANOVA F and p-values are reported in the legends to the Supplementary Figures; significant results of the contrast tests are further indicated by asterisks in the graphs. All statistical tests were two-tailed and $p < 0.05$ was considered statistically significant. For the complete set of raw data, see the data source file.

**Supplementary Figure S1.**

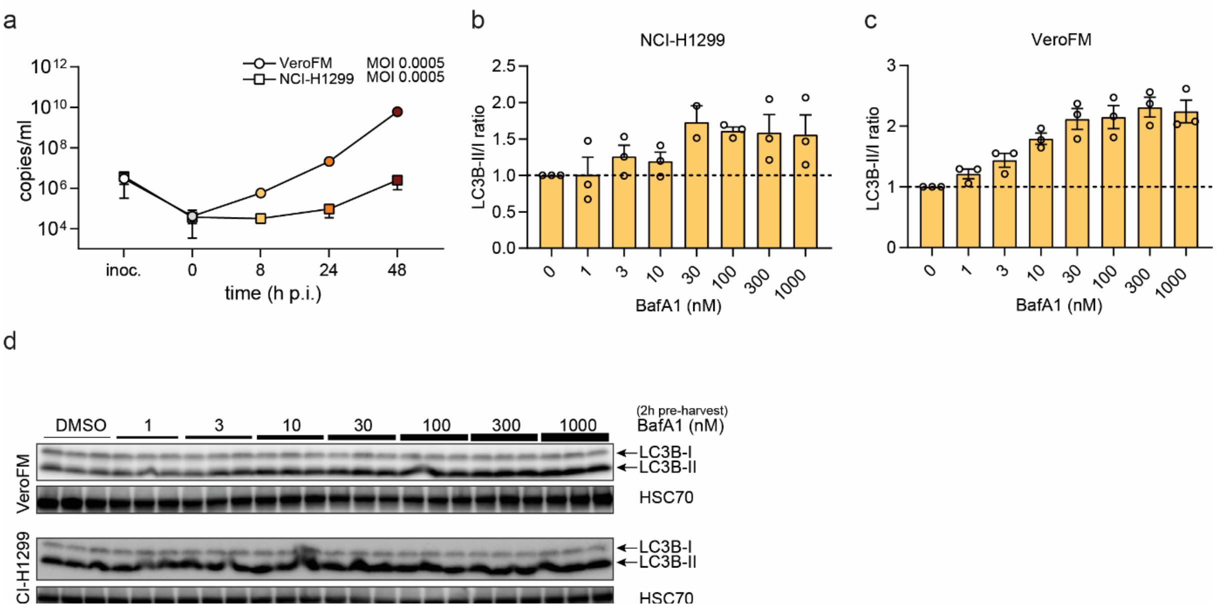


**Fig.S1:** (**a**) Growth of SARS-CoV-2 in NCI-H1299 cells and VeroFM cells over time. NCI-H1299 cells (squares) or VeroFM cells (circles) were infected with SARS-CoV-2 (MOI = 0.0005) and SARS-CoV-2 viral copies per ml were determined by real-time RT-PCR at 8h, 24 h, and 48 h p.i.. (**b-d**) Titration of the BafA1 effect on LC3B lipidation in NCI-H1299 (b,d) cells or VeroFM (c,d) cells as required for the flux assays. Cells were exposed to increasing concentrations of BafA1 for 2 h before cells were harvested and lysed. Ratios of LC3B-II/I were analyzed by Western blotting – representative blots shown in d. Based on these data, 100 nM BafA1 was chosen for the experiments assessing autophagic flux as the concentration achieving complete block of autophagosome-lysosome fusion.

**Supplementary Figure S2.**

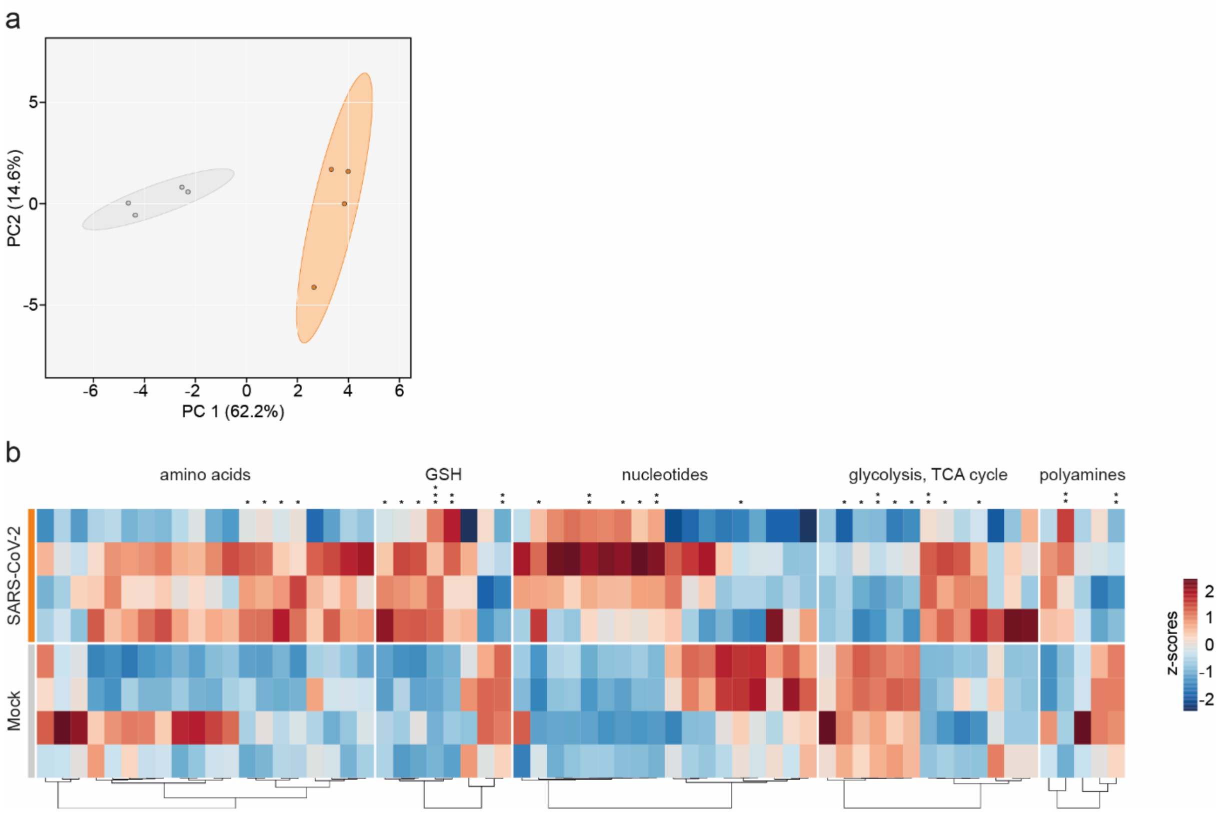


**Fig.S2:** (**a**) Multivariate unsupervised principal component analysis (PCA) comparing mock and SARS-CoV-2 infected (24 h p.i) VeroFM cells using all quantified metabolites for each treatment group. N = 4 per group. (**b**) Heat map with the log2 transformed and z-scored intensities of Mock and SARS-CoV-2 infected (24 h p.i) VeroFM cells for 'amino acids', 'GSH metabolism', 'nucleotides', 'glycolysis/ TCA cycle' and 'polyamine metabolism'. *Adjusted p-value false discovery rate (FDR) ≤ 0.05; **FDR ≤ 0.01; ***FDR ≤ 0.001. N = 4 per group.

**Supplementary Figure S3.**

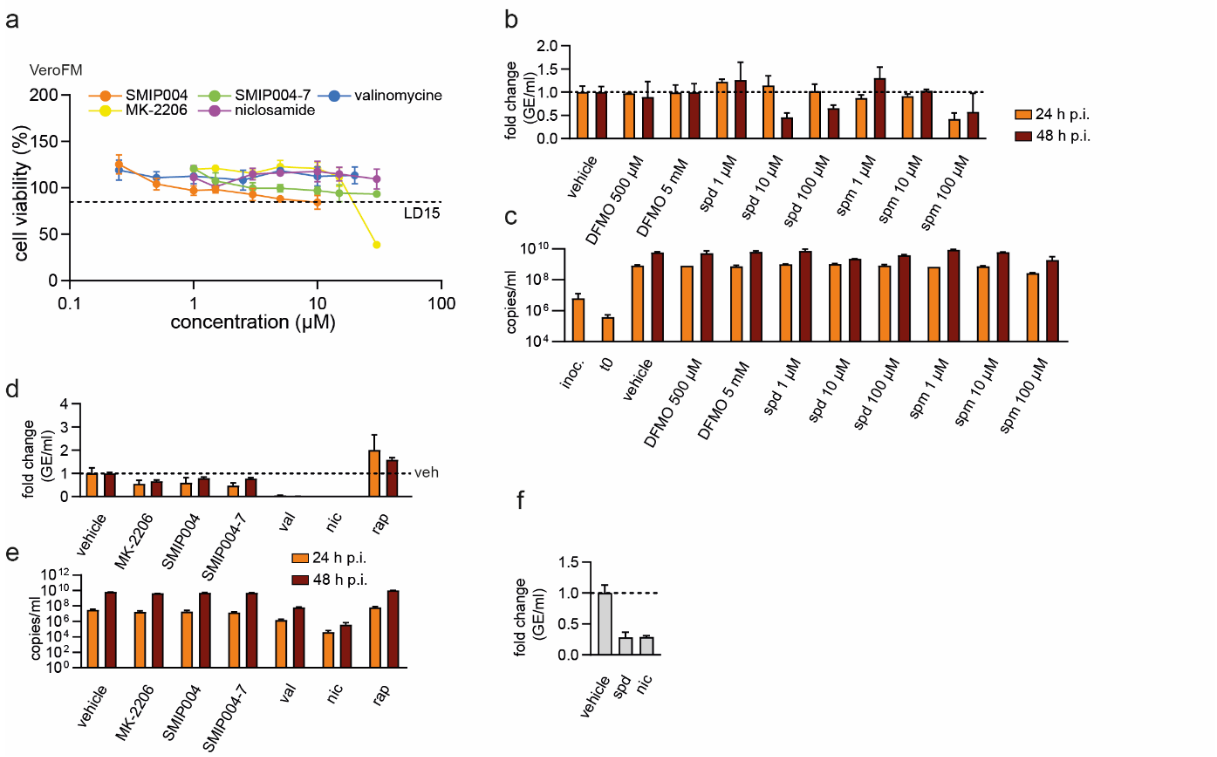


**Fig.S3:** (**a**) Toxicity assays of autophagy modulators. VeroFM cells were treated for 24 h with inhibitors MK-2206, SMIP004, SMIP004-7, valinomycin or niclosamide with increasing concentrations as indicated and cell viability was determined by the MTT (tetrazole 3-(4,5-dimethylthiazol-2-yl-)-2,5-diphenyltetrazolium bromide) assay as described before (*6*). Based on these results, the concentrations used in the experiments were chosen (at least 85% cell viability). (**b,c**) concentration-dependent inhibition of SARS-CoV-2 replication by DFMO, spd (spermidine), spm (spermine) or vehicle. VeroFM cells were infected with SARS-CoV-2 (MOI = 0.0005) and treated with indicated concentrations. SARS-CoV-2 genome equivalents per ml (GE) were determined by real-time RT-PCR at 24 h and 48 h p.i., data are presented as fold change (b) and copies per ml (c). (**d,e**) VeroFM cells were infected analog to b,c and treated with MK-2206 (1 µM), SMIP004 (10 µM), SMIP004-7 (10 µM), val (valinomycin, 5 µM), nic (niclosamide, 10 µM), rap (rapamycin, 300 nM), or vehicle. Real-time RT-PCR data to determine viral genome equivalents are presented as fold change (d) and copies per ml (e). (**f**) VeroFM cells were treated with spd (100 µM), nic (5 µM) or veh (vehicle) 24 h prior to infection with SARS-CoV-2 (MOI = 0.05). SARS-CoV-2 genome equivalents per ml (GE) were determined by real-time RT-PCR at 24 h and 48 h p.i., data are presented as fold change. In all panels error bars denote standard error of mean derived from n = 3 biologically independent experiments.

#### Supplementary Table 1

| Metabolite | HMDB | log2FC (SARS2-control) | p-value | FDR | -lOG10(FDR) |
|---|---|---|---|---|---|
| Cysteinylglycine | HMDB0000078 | 3,19 | 0,000001 | 0,0001 | 4,02 |
| N-Acetylputrescine | HMDB0002064 | 1,65 | 0,000018 | 0,0006 | 3,23 |
| 3-Phosphoglyceric acid | HMDB0000807 | 0,98 | 0,000186 | 0,0041 | 2,39 |
| Putrescine | HMDB0001414 | 1,25 | 0,000708 | 0,0091 | 2,04 |
| Fructose 6-phosphate | HMDB0000124 | -1,26 | 0,000918 | 0,0091 | 2,04 |
| deoxyCTP (dCTP) | HMDB0000998 | 0,65 | 0,001017 | 0,0091 | 2,04 |
| Adenosine | HMDB0000050 | 1,20 | 0,001103 | 0,0091 | 2,04 |
| Homocysteine | HMDB0000742 | 1,56 | 0,001109 | 0,0091 | 2,04 |
| Thymidine 5'-triphosphate (dTTP) | HMDB0001342 | 0,48 | 0,001307 | 0,0096 | 2,02 |
| Spermine | HMDB0001256 | -0,98 | 0,001467 | 0,0097 | 2,01 |
| Uridine triphosphate (UTP) | HMDB0000285 | 0,56 | 0,001765 | 0,0106 | 1,98 |
| L-Lactic acid | HMDB0000190 | 0,41 | 0,002288 | 0,0118 | 1,93 |
| Cytidine triphosphate (CTP) | HMDB0000082 | 0,87 | 0,002332 | 0,0118 | 1,93 |
| cis-Aconitic acid | HMDB0000072 | -0,51 | 0,003456 | 0,0151 | 1,82 |
| Isocitric acid | HMDB0000193 | -0,52 | 0,003655 | 0,0151 | 1,82 |
| L-Glutamic acid | HMDB0000148 | 0,19 | 0,003667 | 0,0151 | 1,82 |
| Glycine | HMDB0000123 | 0,70 | 0,005188 | 0,0201 | 1,70 |
| Guanosine triphosphate (GTP) | HMDB0001273 | 0,42 | 0,005725 | 0,0204 | 1,69 |
| Fructose 1,6-bisphosphate | HMDB0001058 | -0,65 | 0,005873 | 0,0204 | 1,69 |
| Adenosine triphosphate (ATP) | HMDB0000538 | 0,35 | 0,007814 | 0,0258 | 1,59 |
| Oxoglutaric acid | HMDB0000208 | -0,64 | 0,008597 | 0,0270 | 1,57 |
| L-Cysteine | HMDB0000574 | 1,51 | 0,009448 | 0,0283 | 1,55 |
| Adenosine monophosphate (AMP) | HMDB0000045 | -0,65 | 0,009934 | 0,0285 | 1,55 |
| Succinic acid | HMDB0000254 | 0,37 | 0,014020 | 0,0386 | 1,41 |
| L-Tyrosine | HMDB0000158 | 0,96 | 0,016098 | 0,0425 | 1,37 |

**Abbreviations:** NA, not applicable; FC, Fold Change; FDR, False Discovery Rate; HMDB, Human Metabolome Database

#### Supplementary Table 2

| Treatment | MOCK_01 | MOCK_02 | MOCK_03 | MOCK_04 | SARS-CoV-2_01 | SARS-CoV-2_02 | SARS-CoV-2_03 | SARS-CoV-2_04 |
|---|---|---|---|---|---|---|---|---|
| Label | 1 | 1 | 1 | 1 | 0 | 0 | 0 | 0 |
| 3-Phosphoglyceric acid | 791613,101 | 706044,651 | 718039,159 | 733297,265 | 1437379,72 | 1597798,05 | 1615804,05 | 1152595,07 |
| Aconitic acid | 22484,1246 | 20627,5949 | 24264,5436 | 22982,5579 | 16189,7116 | 15568,7688 | 18560,0005 | 13303,373 |
| ADP | 1636091,8 | 1596942,92 | 1891581,25 | 1916390,85 | 1631747,03 | 1726314,24 | 1972885,89 | 1450077,85 |
| alpha-ketoglutaric acid | 57962,8479 | 62006,0295 | 60662,3736 | 60467,7898 | 42160,9665 | 42703,8736 | 43077,6448 | 26317,8204 |
| AMP | 549677,195 | 588526,723 | 754738,12 | 754163,483 | 368159,85 | 472103,242 | 499775,721 | 350100,436 |
| ATP | 3105519,36 | 2918450,19 | 3068063,25 | 3194890,04 | 3516624,76 | 3862246,92 | 4603198,91 | 3711473,3 |
| cAMP | 2301,99667 | 2013,89715 | 2287,87084 | 2639,79499 | 3237,52738 | 1868,62449 | 2143,79845 | 1401,67131 |
| CDP | 257166,818 | 228509,551 | 262388,679 | 272286,499 | 268715,049 | 283316,193 | 286963,146 | 215213,577 |
| Citric acid | 875611,245 | 718591,379 | 783107,152 | 748087,923 | 925105,306 | 673626,203 | 674453,379 | 476724,757 |
| CMP | 54505,9729 | 59763,9227 | 80671,6077 | 72536,4538 | 58012,4764 | 51974,6388 | 48502,8577 | 36690,7159 |
| CTP | 196637,346 | 138848,984 | 118507,599 | 130811,947 | 301643,322 | 251718,637 | 278989,112 | 237066,476 |
| dATP | 26535,8277 | 26232,2889 | 27612,4925 | 29046,1043 | 28945,0013 | 35462,5456 | 43767,4563 | 37085,2895 |
| dCTP | 9637,40207 | 8474,37455 | 8209,32283 | 8704,84294 | 11856,7808 | 13008,1139 | 16281,9938 | 13906,5393 |
| dGTP | 7528,48434 | 7150,38933 | 7591,44228 | 7913,06239 | 7590,08756 | 8841,09274 | 11215,5499 | 9857,47799 |
| dTMP | 15802,6418 | 16187,1903 | 22428,8075 | 21770,4261 | 11868,1855 | 14693,7842 | 15832,0601 | 12924,2198 |
| dTTP | 24430,6094 | 23260,6737 | 24656,8057 | 24558,5402 | 30172,8322 | 32853,5475 | 39286,8755 | 32534,5215 |
| Fructose-1,6-BP | 2294938,58 | 2453673,07 | 2820005,07 | 2884934,75 | 1330858,03 | 1562180,29 | 2106053,8 | 1669834,02 |
| Fructose-6-P | 36504,9613 | 37979,0435 | 42036,4262 | 43047,8097 | 12385,4769 | 13423,4525 | 23517,5797 | 17154,1324 |
| Fumaric Acid | 67244,8051 | 55182,7179 | 55727,8203 | 53224,8282 | 102777,632 | 70283,4455 | 71890,2082 | 53971,6499 |
| GDP | 291233,099 | 306359,974 | 366557,396 | 373581,747 | 273628,713 | 281341,198 | 322829,517 | 248620,847 |
| GTP | 680150,928 | 605540,091 | 637985,43 | 661198,06 | 748473,384 | 825481,179 | 1016542,1 | 877938,829 |
| IMP | 9594,38483 | 13626,8764 | 10502,5121 | 10832,8031 | 10900,5985 | 12917,9926 | 14495,8346 | 11182,599 |
| Isocitric acid | 20697,2388 | 20648,4234 | 23140,5795 | 22609,0883 | 13627,7829 | 15421,5835 | 18395,6165 | 13325,185 |
| Lactic acid | 56635,5002 | 50473,8584 | 55651,0756 | 53927,6068 | 75832,8034 | 70386,422 | 78238,5015 | 62787,674 |
| Malic acid | 48358,6657 | 43492,5722 | 54774,065 | 48498,6512 | 59339,1908 | 59863,5926 | 62057,5403 | 50266,8302 |
| Phosphoenolpyruvate | 37475,6826 | 30768,059 | 29776,4899 | 30216,9835 | 56136,359 | 30837,2391 | 37114,111 | 42473,5725 |
| Pyruvic acid | 569,397773 | 1514,92901 | 574,014559 | 528,905549 | 209,235962 | 179,189583 | 368,971777 | 209,078973 |
| Sedoheptulose-7-P | 435,34535 | 517,103823 | 488,338321 | 385,883165 | 490,800374 | 705,318527 | 777,84746 | 778,353406 |
| Succinic acid | 33569,5583 | 30406,7985 | 37244,4883 | 33879,8189 | 49701,8933 | 44949,5059 | 42663,5559 | 37099,3776 |
| UDP | 978724,421 | 968333,48 | 1115375,05 | 1161477,53 | 932731,546 | 1085042,45 | 1231502,81 | 914168,529 |
| UMP | 175320,877 | 169430,332 | 196251,096 | 180777,676 | 182019,864 | 137678,211 | 137515,404 | 94241,0976 |
| UTP | 533390,212 | 486788,441 | 494723,324 | 540326,274 | 657246,648 | 706202,611 | 896557,782 | 776730,286 |
| 4-Hydroxyproline | 366495,449 | 332364,789 | 334167,236 | 354943,982 | 450521,432 | 388002,268 | 390815,624 | 281465,655 |
| Adenosine | 33769,1354 | 53345,8146 | 36013,7266 | 41046,989 | 82739,732 | 119023,952 | 98863,411 | 75925,8047 |
| Alanine | 6106969,1 | 5898420,2 | 6502838,92 | 5855731,17 | 6327098,69 | 6417773,25 | 6736962,3 | 5322586,47 |
| Arginine | 2693888,17 | 5753651,18 | 2755585,83 | 2720035,09 | 2752327,98 | 2366267,66 | 2931425,85 | 2418402,42 |
| Asparagine | 1737224,44 | 2853292,93 | 1872148 | 1773546,06 | 2459853,85 | 2159524,34 | 2429967,15 | 1973586,92 |
| Aspartic Acid | 2458476,94 | 2524517,33 | 2684334,73 | 2519013,32 | 2982351,9 | 2963323,18 | 3277727,59 | 2574165,98 |
| beta-Alanine | 29416358,9 | 27234150,3 | 28711363,9 | 30055026,8 | 27284130,9 | 24373927,2 | 27595087,7 | 19951736,7 |
| Carnosine | 17622,7756 | 16549,8103 | 17093,0337 | 16730,8887 | 18630,5846 | 14015,0302 | 23398,0135 | 14463,961 |
| Citrulline | 54475,643 | 52422,7245 | 59006,9254 | 51809,4552 | 59454,7411 | 58339,0729 | 72726,3332 | 44744,33 |
| Creatine | 590655,498 | 424574,405 | 210259,935 | 361289,806 | 567289,583 | 347548,633 | 445210,705 | 263926,346 |
| Cystathionine | 226615,015 | 236177,507 | 242506,843 | 199319,345 | 281102,176 | 256252,728 | 341458,191 | 243281,886 |
| Cysteine | 53559,3196 | 55706,5627 | 25447,787 | 27932,2547 | 172079,923 | 124029,726 | 98739,6266 | 69649,1993 |

| | | | | | | | | |
|---|---|---|---|---|---|---|---|---|
| Cysteinyl-Glycine | 111784,924 | 101956,278 | 77571,9214 | 85516,4146 | 947525,977 | 937663,703 | 666403,405 | 893891,431 |
| gama-aminobutyric acid | 22494302,1 | 21384053,7 | 21486388 | 19453555,6 | 21971760 | 20327050,3 | 23512866 | 16391917,5 |
| Glutamic Acid | 58219049,2 | 60752887,1 | 61401800,4 | 57723763,7 | 70050872,8 | 68259310,7 | 70828071,6 | 63414239,4 |
| Glutamine | 25496509 | 27696470,5 | 26228488,5 | 27416950,7 | 25206113,8 | 24694803,5 | 26764148,3 | 24179458,9 |
| Glutathione | 16270113 | 13889746,4 | 16352641,7 | 14897398 | 15929979,3 | 15330938,6 | 16021120,8 | 11613259,1 |
| Glycine | 364853,024 | 500012,235 | 358392,045 | 327483,419 | 687794,883 | 615314,042 | 694637,703 | 518815,737 |
| Histidine | 2309219,75 | 2525595 | 2077361,6 | 1813544,06 | 2432188,72 | 2294149,19 | 2457117,74 | 1981290,78 |
| Homocysteine | 5585,10299 | 3127,24065 | 3303,56447 | 5434,97398 | 11757,8628 | 17119,0849 | 12360,6299 | 10390,2371 |
| Hypotaurine | 5303323,47 | 5191700,08 | 6176338,21 | 5790835,09 | 5333552,68 | 5484303,11 | 5638761,48 | 4338015,47 |
| Isoleucine | 3681540,41 | 3493794,39 | 2869801,48 | 2752341,08 | 3882390,71 | 3486985,23 | 3417948,12 | 3112794,03 |
| Leucine | 2850541,63 | 3311258,06 | 2649611,28 | 2546906,35 | 3354294,29 | 3025504,52 | 3274648,76 | 2690583,11 |
| Lysine | 7020120,19 | 8676890 | 7119269,99 | 6976380,49 | 6427459,72 | 7313868,71 | 6779176,06 | 6119384,23 |
| Methionine | 722863,304 | 847127,974 | 615480,232 | 616220,72 | 806052,342 | 860695,875 | 845828,491 | 706214,661 |
| Methylthioadenosine | 59179,3049 | 59218,7973 | 19043,4466 | 26249,4804 | 90627,4102 | 91964,3884 | 75816,0909 | 57001,7578 |
| N-acetylputrescine | 77001,1038 | 97089,0719 | 82331,2982 | 80653,2576 | 237138,427 | 291359,135 | 304632,998 | 221229,936 |
| Phenylalanine | 2024086,6 | 2191846,85 | 1856223,98 | 1797035,7 | 2449138,88 | 2185111,94 | 2356421,35 | 1859079,06 |
| Proline | 12534703,8 | 10888717,3 | 12433863,3 | 11907719,8 | 15483644 | 13494245,9 | 15829387,7 | 10662380,1 |
| Putresince | 6515042,38 | 9213629,63 | 7912005,98 | 7634479,49 | 16163468,7 | 14229054,3 | 20636028,8 | 23572926,6 |
| Serine | 635438,302 | 953888,118 | 608405,618 | 582586,461 | 776110,058 | 754775,657 | 804411,971 | 714270,618 |
| Spermidine | 6542075,59 | 9128438,21 | 9127046,02 | 8100036,36 | 4245190,77 | 3119061,02 | 6283349,83 | 7041080,77 |
| Spermine | 1174893,96 | 1446179,17 | 1488899,87 | 1502433,97 | 710601,187 | 527812,587 | 909577,91 | 691756,656 |
| Taurine | 24836921,1 | 22903726,6 | 28863962,3 | 24214828,7 | 23355737,5 | 24274491,2 | 27192006,2 | 19170388 |
| Threonine | 2229980,6 | 2369546,07 | 2334098,71 | 2081583,13 | 2687675,09 | 2443021,1 | 3037515,65 | 2173279,66 |
| Tryptophane | 940383,067 | 1146624,23 | 854009,065 | 873019,929 | 1053985,26 | 964745,669 | 1085245,29 | 854257,531 |
| Tyrosine | 1447095,98 | 1731791,45 | 1083476,44 | 802851,585 | 2821275,71 | 3098989,93 | 2032496,77 | 1905071,56 |
| Valine | 3508459,34 | 4145672,64 | 3330156,32 | 3263133,12 | 3870254,9 | 3396594,51 | 4254311,44 | 3062814,19 |

248

References (Supplementary Material)
